# Sexual conflict over mating duration and frequency in *Zygogramma bicolorata*

**DOI:** 10.1101/2025.02.26.640458

**Authors:** Rabi Sankar Pal, Anirban Bhowmick, Bodhisatta Nandy

**Author notes:** Corresponding author, contact number.

## Abstract

Duration and frequency of mating are the most prominent aspects of mating behaviour over which sexual conflict can occur. Parthenium beetle (*Zygogramma bicolorata*) exhibits an extremely high mating rate, with each mating typically lasting several hours. We investigated mating behaviour in this beetle to assess if there is sexual conflict. Theories of sexual conflict predict that such mating behaviour, while being beneficial to the males, should be detrimental to the females with regard to reproduction and/or survival. We experimentally manipulated the length and number of matings, and examined the effect on fecundity and survivorship of the females. While long mating has previously been identified as a mate-guarding strategy that benefits the males, our data reveal that it is detrimental to female fitness, supporting the sexual conflict hypothesis. Further, we checked natural variation in mating duration and its correlation with key fitness components. Our results show that female longevity is negatively correlated to reproductive rate, but positively correlated to lifetime reproduction. We argue that such a pattern is consistent with sexual conflict. While being the first clear demonstration of sexual conflict in this species, we discuss its significance, including a potential role in reducing its efficacy as a biocontrol.

## Introduction

In absence of strict monogamy, fitness interests of sexes often do not coincide, resulting in sexual conflict (Kokko & Jennisons, 2014; Parker, 2006; but see Hosken et al., 2009). Under such conflict, expression of a trait may maximise fitness in one sex while reducing the same in the other (Chapman et al., 1995; Parker, 1979; Rice, 1996). Species with unique mating behaviour, including behavioural extremes, can often offer opportunities to test the predictions of sexual conflict theory, including its putative roles in important ecological processes. One instance of such behavioural extreme is the species with extremely long mating, where long mating is generally considered an example of post-copulatory or copulatory mate guarding (Linn et al., 2007; Pal et al., 2025; Schöfl, G., & Taborsky, 2002). However, little is known about the sexually antagonistic consequences of such long mating, and its cascading ecological consequences.

Mating involves several costs that are functions of time – *viz.*, energy expenditure during mating (Daly, 1978; Watson, 1998), risk of injury or infections (Crudgington & Siva-Jothy, 2000; Hurst et al., 1995; Reinhardt et al., 2015), receipt of harmful substances (Mueller et al., 2007; Wolfner, 2002), increased risk of predation (Magnhagen, 1991; Wing, 1988), lost foraging (Abrahams, 1993; Rowe, 1992) and mating opportunities (Alcock, 1994; Hasselquist & Bensch, 1991; Parker, 1970). For females, the purpose of mating is accomplished once they receive a sufficient number of viable sperm and other resources provided by the males (South & Lewis, 2011). As a result, the gain curve of the females tends to saturate very quickly after the initiation of mating (Arnqvist & Rowe, 2005). Shorter mating durations are, hence, expected to be favoured in females, especially when sperm transfer happens within a short duration (Arnqvist & Rowe, 2005). Mating duration, on the contrary, is an important determinant of sperm competition success in males (Simmons, 2002). Besides sperm transfer, male investment in seminal fluid often increases their fertilisation success (Kelly & Jennions, 2011; Wigby et al., 2009). The male gain curve, therefore, saturates at a longer time compared to the female’s, resulting in a longer optimal mating duration in males (Arnqvist & Rowe, 2005).

Males may manipulate females by inducing nonreceptivity, thereby taking the female mating rate below the optimum (Manning, 1967; Nallasivan, 2024; Ravi Ram et al., 2005), or seduce/coerce/entice females to mate beyond their optimum (Arnqvist & Nilsson, 2000; Lessells, 2006). Longer mating may work in favour of the males if it ensures nonreceptivity or reduced receptivity to future matings, even if for a finite period (Chapman et al., 2003; Chen et al., 1988; Wedell, 2005; Yang et al., 2023). Mating in Parthenium beetle *Zygogramma bicolorata* continues for several hours that involves repeated intromissions (Afaq & Omkar, 2017; Bhaisare et al., 2021; Bhaisare et al., 2024; Pal et al., 2025) though sperm transfer takes place during the early phase of such mating and requires only a brief period (Pal et al., 2025). There could be asymmetry in the cost paid by males and females due to such long mating (Mazzi et al., 2009; Rowe, 1994). *Zygogramma bicolorata* females carry the males during the entire mating, which may interfere with foraging (e.g. Rowe et al., 1994). Interestingly, females show male-repelling behaviours, e.g. kicking and shaking (Pal et al., 2024), suggesting that such extended mating may not be favourable to females. Further, long mating can modulate female remating behaviour, potentially reducing female fitness (Pal et al., 2025). The cost incurred by the males during mating may come from the production and transfer of ejaculate, performing possible copulatory courtship behaviours, and starvation during the long mating period (Pal et al., 2025). On the other hand, the cost of such long mating and repeated intromissions on females is not known.

In this study, we investigated the scope of sexual conflict surrounding the unique mating behaviour in *Z. bicolorata*. To detect any subtle signal of sexual conflict, we experimentally manipulated the duration of mating and the number of mating opportunities for the females, and looked into the potential consequences on their fitness. If there is sexual conflict, long mating and higher mating rates should be more detrimental to females than short and infrequent mating. We then looked at a set of unmanipulated matings, and assessed pairwise correlations between a set of mating behaviour and fitness components. This was done to assess if natural variation in mating behaviour translates into detrimental fitness effects in a manner predicted by sexual conflict over mating rate and duration.

## Methods

*Zygogramma bicolorata* Pallister is a herbivorous beetle of the family Chrysomelidae. These beetles are specialist feeders and a biocontrol agent of the weed Parthenium (Adkins et al., 2018). With little sexual dimorphism, promiscuous mating system, and intriguing mating behaviour, this species is an interesting system to study sexual conflict. Details on the life attributes of these beetles can be found in Patel et al. (2020).

Present study was conducted on a laboratory population of these beetles (establishment and laboratory maintenance can be found in Pal et al. 2024). Briefly, a lab population of 350–400 individuals per generation was maintained at laboratory conditions (26 ± 2 °C, 24-hour light). Larvae were grown in the aerated Petri dishes (90mm diameter, 15 mm height) in controlled density with *ad libitum* Parthenium leaves. Fourth instar larvae were transferred to moist sand to facilitate pupation. Adults were held in groups of 10 in single-sex in transparent containers (adult boxes) with *ad libitum* Parthenium leaves upon eclosion. The assays commenced on day 11 post-eclosion when they reached sexual maturity. Experiment 1 was conducted in the 11th laboratory generation, whereas Experiment 2 was conducted in the first laboratory generation to capture the natural variation in the traits.

### Experiment 1

In the first experiment, we aimed to test the predictions from sexual conflict theory. If there is sexual conflict over mating in the Parthenium beetle, longer and more frequent matings should be detrimental to the females compared to shorter and less frequent mating. Unmated females and males were randomly paired into a petri dish with Parthenium leaf (mating arena) on post-eclosion day 11. They were allowed either short (interrupted after 2 hours of mating, or complete mating of ≤ 2 hours, approximately 30 intromissions) or complete, long (uninterrupted) mating. The males were removed following interruption or completion of the mating and held in groups of 10 in the adult boxes. The females from each of the short and long mating durations were further divided into two groups randomly. The first group was allowed no further exposure to males and was held individually in Petri dish with fresh Parthenium leaf provided on every alternate day. The females in the second group were released in the mating arena on every alternate day with randomly picked males (that mated previously) until 15 such exposures. The females from short or complete mating treatments were allowed the same duration of mating from the initiation of intromission on every successful mating exposure. This generated four combinations, *viz.,* Single-short (SS) (n=32), multiple-short (MS) (n=35), single-complete (SC) (n=35), and multiple-complete (MC) (n=33) mating (Figure S1). Following every male-female exposure/mating, males were released back into the adult boxes, while females were held individually in the Petri dishes with freshly supplied leaves. Pairs that did not mate within four hours since the release into the mating arena were removed and kept in their respective places. The fecundity of the females with individual IDs was measured every alternate day till day 31 since their first mating (48 hours after the last male-female interaction). MS and MC females mated 13.1 and 12.47 times on average (Mean ± SE: MS 13.1 ± 0.393; MC 12.467 ± 0.317) out of 15 exposures to males.

Following this, the males were discarded, and the females were kept in empty Petri dishes for starvation. Some females died before reaching day 31. Fecundity data from these females were excluded from the analysis. At the beginning of the starvation assay, SS, MS, SC, and MC combinations had 29, 30, 29, and 30 females, respectively. The females were checked for mortality every day at the same time until all the females died.

Pal et al. (2025) found no difference in sperm transfer between short mating of 30 intromissions (∼2 hours) and uninterrupted mating. Hence, any difference in fecundity is unlikely to be confounded by the number of sperm transferred to the females. Cumulative early-life fecundity of the females was checked till 31 days (∼33% of their lifespan, which contributes to more than 40% of the lifetime reproductive output of the females: see Pal et al., 2025). Reproductive ageing seems to set in beyond day 20 post-mating (Pal et al., 2025).

### Experiment 2

In a natural population of the beetle, mating duration varies substantially (Pal et al., 2024). Hence, it is important to assess if such variation correlates to variation in key fitness components, especially those that can be tied to sexual conflict. Therefore, we designed a second experiment to measure the degree of correlations between different components of fitness (longevity, lifetime reproductive output, reproductive rate), and mating behaviour (mating duration, number of intromission). Since the degree of sexual conflict is often affected by body size variation, we also included it in the investigation. If there is sexual conflict, negative correlation between some of the fitness components, and mating duration or number of intromissions should be observed.

On day 11 post-eclosion, individual beetles were assigned an identification number, immobilised in mild CO_2_ anaesthesia, and dorsal image of the beetles was captured in Zeiss Stemi 508 microscope fitted with Zeiss Axiocam ERc5s using Zen 2.3 (blue edition) software. The length of the left elytron (usually a proxy of the body size in beetles; Baar et al., 2018; Matsumoto & Knell, 2017) along the elytral suture was measured accurate to 0.001 μm. Following measurement, beetles were kept individually in glass vials (25 mm diameter, 96 mm height) with leaf for 24 hours for pre-conditioning prior to the observation. Following pre-conditioning, males and females were randomly paired in glass vials with small pieces of Parthenium leaves. The long mating of *Z. bicolorata* consists of a series of repeated intromissions (see Pal et al., 2025). The mating duration and the number of intromissions within a single mating were observed for each of the 48 pairs closest to a minute. Males were removed immediately upon completion of mating, and females were kept individually in Petri dishes with fresh Parthenium leaves, which were replaced daily. The fecundity of individual females was measured every day at a fixed time until all the females died. Of the initial 48 females, one female was lost due to handling error. The final sample size for this part of the assay was 47. Pairwise correlations were checked between female body size, male body size, mating duration (the time between mounting and dismounting of a pair that included repeated intromissions), number of intromissions (total number of intromissions in a single complete mating), longevity (number of days the females survived post-mating), lifetime reproductive output (total number of eggs laid in their lifetime) and reproductive rate (lifetime reproductive output per unit day of lifespan).

### Statistical Analyses

All the analyses were performed in R version 4.2.0 (R Core Team, 2022). The normality of the residuals was checked using the Shapiro-Wilk test. Anova function from the `car` package was used to obtain *p* values for the predictor variables and their interactions (Fox & Weisberg, 2019). In experiment 1, cumulative early-life fecundity was fitted into negative binomial generalised linear models with log link functions using the `MASS` package to correct for overdispersion (Venables & Ripley, 2002). The number of male exposure/mating and the duration of mating were modelled as fixed factors in the analyses. Further, `emmeans` package was used to find pairwise contrasts (Lenth, 2022). The effect of the number of male exposure/mating and the duration of mating on female survival under starvation was analysed taking them as fixed factors in the Cox Proportional Hazard models using the `survival` package (Therneau, 2022). Plots were created using the `ggplot2` package (Wickham, 2016). In experiment 2, females with zero lifetime fecundity (and reproductive rate) were removed prior to the correlation test. Pearson correlation coefficients were computed to check for significant linear association between the variables.

## Results

The results of Experiment 1 showed that the number of mating/male exposures and the duration of mating did not show a significant effect on the cumulative early-life fecundity of the females. However, duration × exposure interaction significantly impacted the cumulative early-life fecundity (Table 1). In the pairwise comparison, only MS and SS groups were statistically different (Figure 1, Table S1). Female survival under starvation was significantly affected by the number of mating/male exposure, and the duration of mating (Table 1). However, the effect of the interaction between these factors was marginally non-significant. Pairwise comparisons revealed significant differences between MC and three other treatment groups (Figure 2, Table S2).

**Figure 1:**
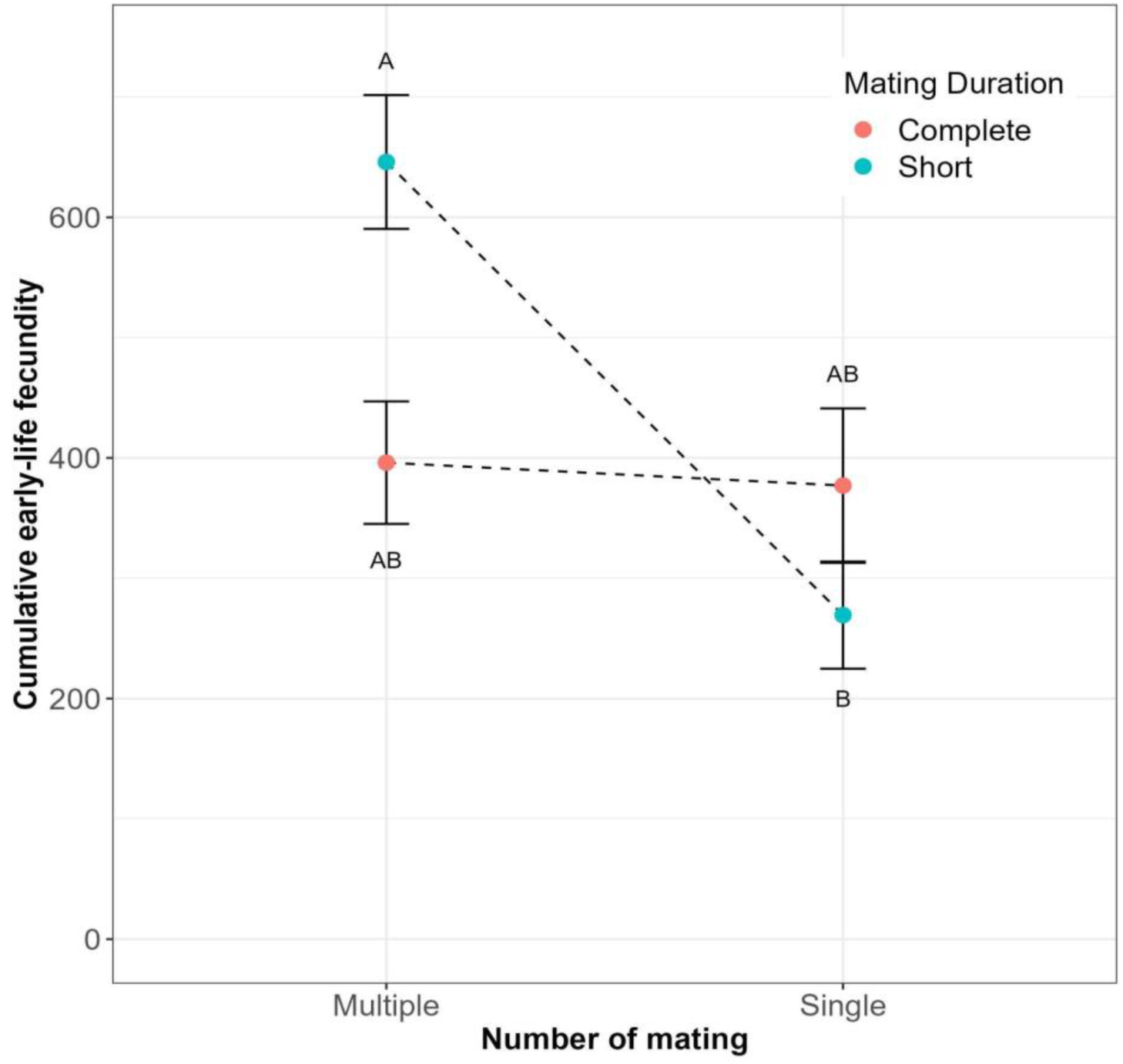
The effect of number of mating/male exposure and mating duration on cumulative early-life fecundity of the females. Treatment groups not sharing the same letter are statistically different.

**Figure 2:**
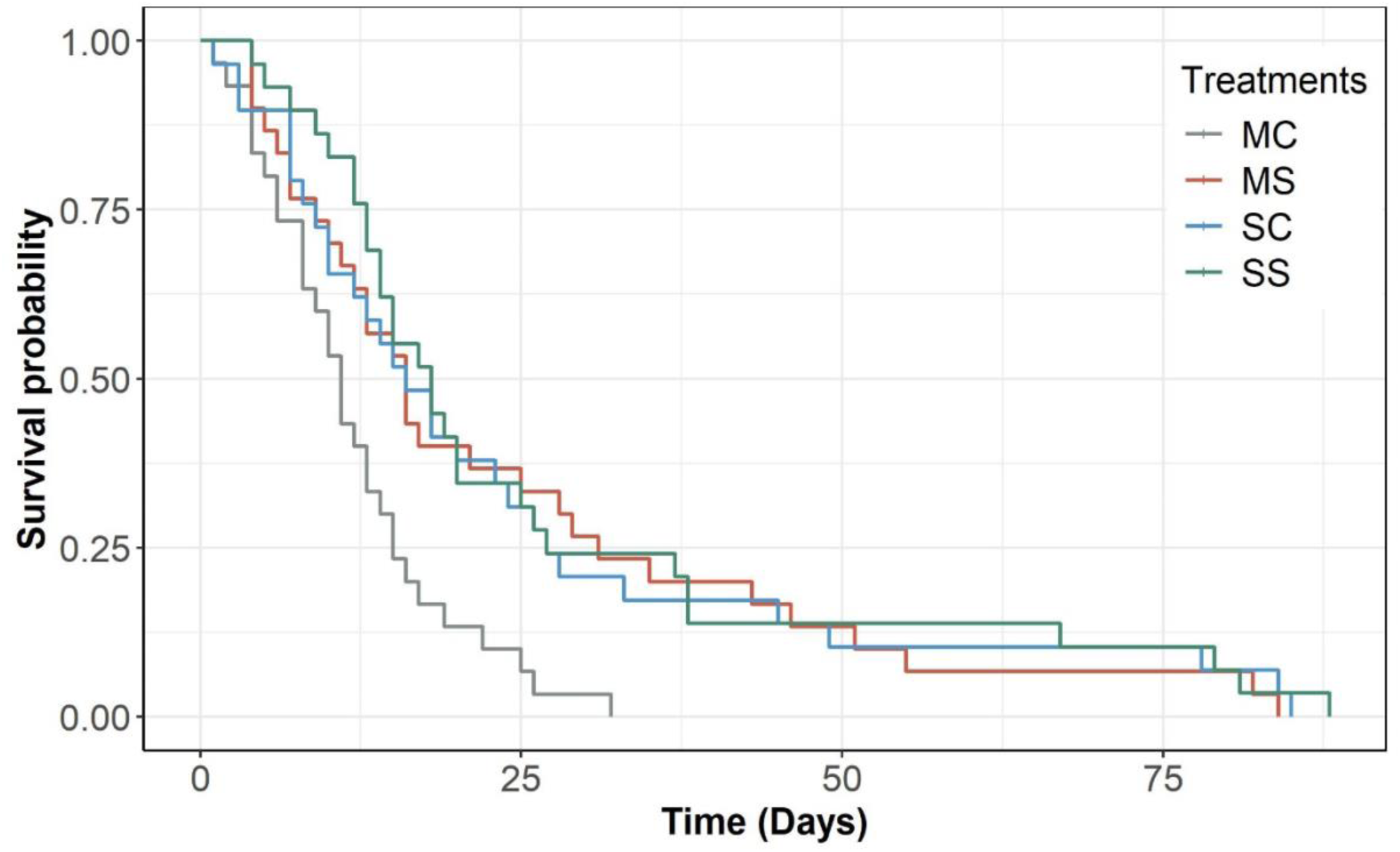
Survival probability of the females of four treatment groups under starvation. The survival probability of MC was significantly different from the other three treatment groups.

**Table 1:**
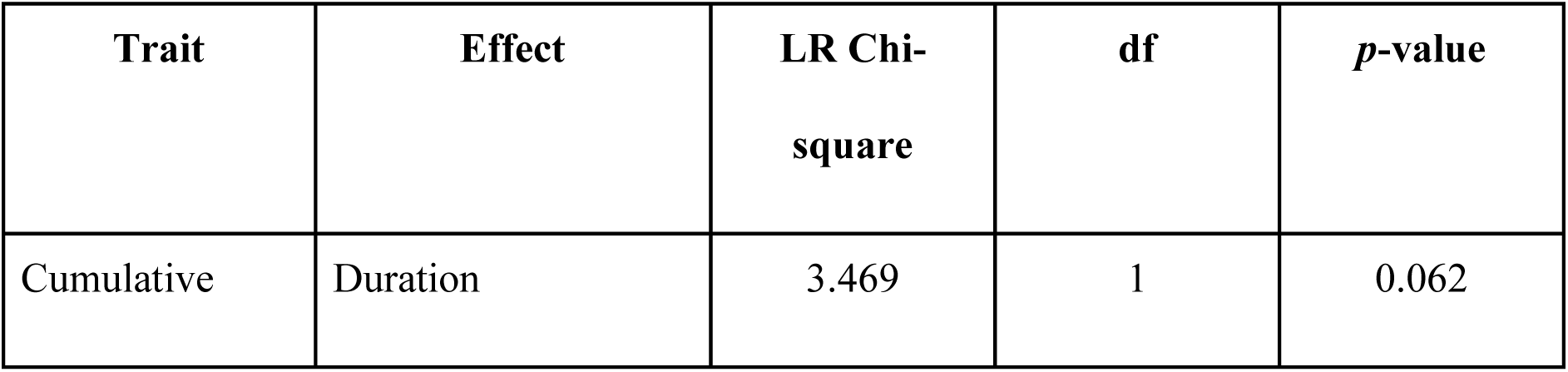
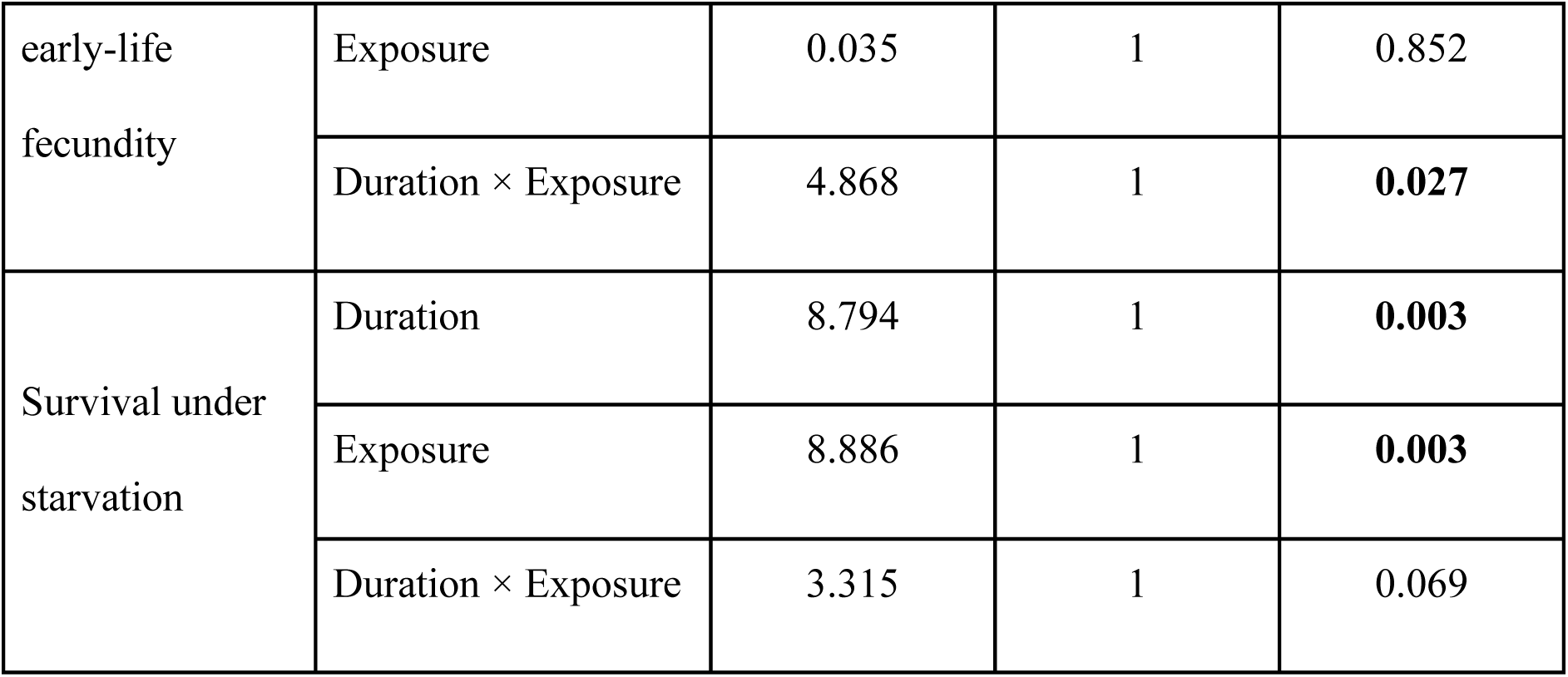
Summary of the results showing the effect of duration and number of mating on cumulative early-life fecundity, and survival under starvation. All the tests were performed considering α= 0.05. Statistically significant *p*-values are highlighted in boldface.

In Experiment 2, a strong positive correlation between the duration of mating and the number of intromissions was understandably evident (*r* = 0.729, *p* <0.001). A positive correlation was also observed between male body size and the number of intromissions (*r* = 0.303, *p* = 0.038). Interestingly, the reproductive rate of the females showed significant positive correlations with the lifetime reproductive output (*r* = 0.405, *p* = 0.007), but a significant negative correlation with the longevity of the females (*r* = −0.554, *p* <0.001). The correlation matrix can be found in Table S3. All other correlation coefficients were not statistically different from zero.

## Discussion

We sought to understand whether the inordinately long mating (consisting of multiple intromissions), along with the high mating rate observed in *Zygogramma bicolorata,* results in sex-specific fitness outcomes. While previous investigations (Afaq & Omkar, 2017; Bhaisare et al., 2024; Pal et al., 2025) showed putative male-benefiting effects, results from our experiments suggest that such mating behaviour is detrimental to female fitness. The best possible scenario for females seems to be multiple short matings, which was found to be associated with the attainment of the highest fecundity, as well as survival rate. Yet, females are known to undergo multiple long matings. Thus, there are clear signs of sexual conflict in this species regarding mating length and number.

In the absence of sexual conflict, multiple complete matings should increase the fitness of both sexes. However, while both mating number and length significantly affected female survival, repeated complete matings (*i.e.*, MC treatment) resulted in significantly higher mortality in females compared to all other treatment groups. When mating was short, cumulative early-life fecundity was significantly lower when only a single mating was allowed, possibly due to sperm limitation. Compared to that, females were found to be almost twice as fecund when multiple matings were allowed. Such an effect of the number of matings was not found when matings were allowed to be complete (as opposed to short/truncated). Interestingly, among all treatment groups, females in the multiple short mating (MS) group appeared to have the highest fecundity. On the other hand, engaging in long mating multiple times did not increase female fitness relative to only a single complete mating. Putting these two lines of observations together, it appears from our results that long mating and high mating rates are indeed detrimental to the females. Since sperm transfer happens in the early phase of mating in this species (Pal et al., 2025), the remaining period may involve the transfer of non-sperm components of the seminal fluid, including accessory gland proteins (Chapman, 2001; Gilchrist & Partridge, 2000; Ravi Ram & Wolfner, 2007). At least some of these seminal fluid proteins can be harmful to females (Chapman et al., 1995; Wigby et al., 2020). These deleterious effects may accumulate over successive matings, reducing female survival.

High cost of mating is predicted to result in decreased mating frequency and increased choosiness in females (Bleu et al., 2012). Surprisingly, females exposed multiple times to males, mated 12–13 times out of 15 mating opportunities. The high frequency of remating in females could represent male coercion (Arnqvist & Nilsson, 2000; Clutton-Brock & Parker, 1995). Whether there is sufficient selection pressure, and existing heritable variation in the relevant traits for female resistance to evolve should be a matter of future investigations.

Interestingly, analysis of cross-correlations between fitness components and behavioural attributes after a single complete mating suggested a positive correlation between the number of intromissions (within a mating) and male body size. If the number of intromissions in a mating is primarily determined by the males, larger males may possibly coerce a higher number of intromissions (Shine & Mason, 2005; Thornhill & Sauer, 1991). Previous report suggests that higher number of intromissions modulates female remating duration, which could be beneficial for males if it results in greater paternity share (Pal et al., 2025). Alternatively, higher number of intromissions with larger males may also represent cryptic female mate choice, especially if body size indicates the quality of the males (Eberhard, 1996; Omkar & Afaq, 2013; Simmons, 1986). However, neither female reproductive rate nor lifetime reproductive output was found to have a positive correlation with the number of intromissions or male size in our investigation. Hence, female interest in higher number of intromissions per mating appears unlikely.

Reproductive rate (eggs produced per day), after single complete mating, was found to have a strong negative correlation (Pearson correlation coefficient = −0.554, *p* < 0.001) with longevity, potentially implying a trade-off between reproduction and longevity (Flatt, 2011; Kirkwood & Rose, 1991; Tatar & Carey, 1995; Van Noordwijk & De Jong, 1986). However, contrary to the predictions from such a trade-off theory, we found a positive correlation, though marginally non-significant, between lifetime reproductive output and longevity (Pearson correlation coefficient = 0.295, *p* = 0.055). Since the longevity–reproduction trade-off is more sensitive to lifetime reproductive investment (Chippindale et al., 1993; Partridge & Fowler, 1992; Dasgupta et al., 2022), there seems to be insufficient evidence to support a trade-off. Interestingly, lifetime reproduction was found to be positively correlated with reproductive rate (Pearson correlation coefficient = 0.405, *p* = 0.007). This indicates that the net fitness impact of higher reproductive rate upon a single complete mating was positive in spite of mortality costs associated with it. Nonetheless, it is reasonable to deduce that the putative effect of reproductive rate on longevity is perhaps independent of lifetime reproductive investment and the possible reproduction–longevity trade-off. In fact, the observed negative correlation between reproductive rate and longevity is more likely to be an indication of mate harm caused by the males, especially if higher reproductive rate is an effect of seminal fluid proteins transferred by the males during the long mating. More importantly, high mating rate (see above) in this species may further offset female fitness by reducing life expectancy. While higher reproductive rate should be in favour of male fitness, it can be significantly detrimental to female fitness, which is often lifetime progeny output (Bonduriansky et al., 2008). However, the latter was not observed in our experiment. Nonetheless, higher reproductive rate early in life may also favour females if female life expectancy is low due to external factors (such as, environmental stresses, predation, diseases etc.) prevailing in natural setups outside the laboratory (Hayward et al., 2015; Stearns et al., 2000; Travers et al., 2015). Therefore, observations on the natural populations of these beetles are essential to test this theory conclusively.

Sexual conflict is known to reduce a population’s average fitness (Arnqvist & Tuda, 2010; Rankin et al., 2011). The fitness lag generated by sexual conflict, often referred to as gender load (Rice et al., 2006), can have far-reaching consequences in natural populations and ecosystems they are part of. For example, if gender load in *Z. bicolorata* results in reduced fecundity and increased mortality in females, a population’s growth rate might be significantly less than theoretically expected. Since the beetle is a specialist herbivore, the regulatory effect of the same on the population of the host plant may be overestimated if intrinsic factor such as the intensity of sexual conflict is not taken into account. Importantly, these beetles were introduced in India as a Biological control for the invasive weed – *Parthenium hysterophorus* in the 1980s and have been surprisingly less effective in its role as a Biocontrol (Adkins & Shabbir, 2014; Maharjan et al., 2020). Future investigations should assess the significance of gender load in these beetle populations in the wild. Generally, the importance of sexual conflict in community/ecosystem-level phenomena needs further investigation.

## Acknowledgements

The authors thank Ranjit Kumar Sahoo for his guidance in establishing the laboratory population of beetles used in this study. We acknowledge the Indian Institute of Science Education and Research Berhampur for financial support in the form of (a) Junior and Senior Research Fellowship to RSP, (b) intramural research grant IG/010218/B0008.

## Notes

### Competing Interest Statement

The authors have declared no competing interest.

