## Supplementary materials for "Sexual conflict over mating duration and frequency in *Zygogramma bicolorata*"

**Supplementary information**

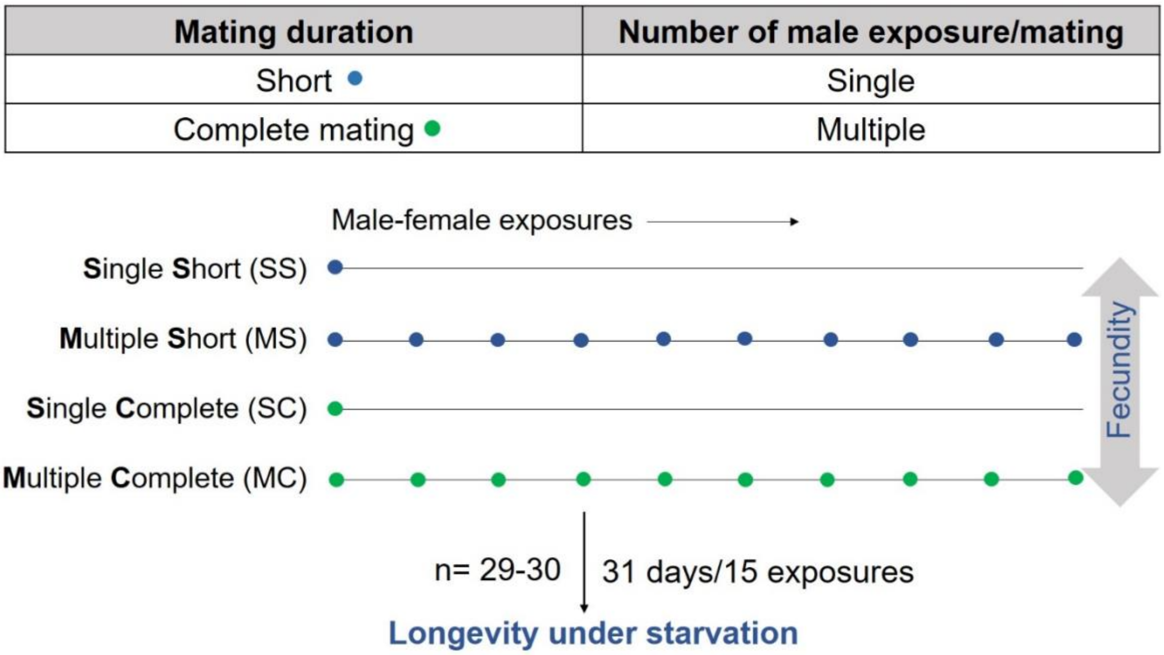


**Figure S1:** Schematic representation of Experiment 1. Treatment groups were allowed single or multiple mating (indicated by closed circles), each of which continued either for a short period (~2 hours; marked in blue) or complete mating (uninterrupted; marked in green). Fecundity was measured till day 31 post-mating, following which females were kept under starvation.

**Table S1:** Results of pairwise comparison between treatments in the cumulative early-life fecundity. The statistically significant *p*-value is highlighted in boldface.

| Contrast | Estimate | SE | df | *z*-ratio | *p*-value |
|---|---|---|---|---|---|
| MC-MS | -0.489 | 0.261 | Inf | -1.872 | 0.24 |
| MC-SC | 0.049 | 0.264 | Inf | 0.187 | 0.998 |
| MC-SS | 0.386 | 0.264 | Inf | 1.462 | 0.46 |

| | | | | | |
|---|---|---|---|---|---|
| MS-SC | 0.538 | 0.264 | Inf | 2.043 | 0.172 |
| MS-SS | 0.875 | 0.264 | Inf | 3.318 | **0.005** |
| SC-SS | 0.336 | 0.266 | Inf | 1.265 | 0.585 |

**Table S2:** Pairwise comparison table showing differences between treatments in their survival under starvation. Statistically significant *p*-values are highlighted in boldface.

| | **MC** | **MS** | **SC** |
|---|---|---|---|
| **MS** | **0.009** | | |
| **SC** | **0.01** | 0.924 | |
| **SS** | **0.003** | 0.814 | 0.814 |

**Table S3:** Correlation matrix showing the results of correlation between the traits. The top-right and the bottom-left off-diagonals of the matrix show the Pearson correlation coefficient (in italics) and the corresponding *p*-values, respectively. Correlation tests were performed considering α= 0.05. Statistically significant *p*-values and the corresponding correlation coefficients are highlighted in boldface.

| | Female body size | Male body size | Duration of mating | Number of intromissions | Longevity | Lifetime reproductive output | Reproductive rate |
|---|---|---|---|---|---|---|---|
| Female body size | × | *-0.09* | *-0.052* | *0.053* | *-0.118* | *-0.139* | *0.047* |
| Male body size | 0.546 | × | *0.198* | ***0.303*** | *-0.099* | *-0.074* | *-0.042* |
| Duration of mating | 0.724 | 0.182 | × | ***0.729*** | *-0.027* | *0.072* | *0.149* |
| Number of intromissions | 0.718 | **0.038** | **<0.001** | × | *-0.146* | *0.057* | *0.257* |
| Longevity | 0.431 | 0.514 | 0.857 | 0.328 | × | *0.295* | ***-0.554*** |
| Lifetime reproductive output | 0.372 | 0.639 | 0.647 | 0.718 | 0.055 | × | ***0.405*** |
| Reproductive rate | 0.763 | 0.787 | 0.341 | 0.096 | **<0.001** | **0.007** | × |
